# Weakly activated core inflammation pathways were identified as a central signaling mechanism contributing to the chronic neurodegeneration in Alzheimer’s disease

**DOI:** 10.1101/2021.08.30.458295

**Authors:** Fuhai Li, Abdallah Eteleeb, William Buchser, Guoqiao Wang, Chengjie Xiong, Philip R. Payne, Eric McDade, Celeste M. Karch, Oscar Harari, Carlos Cruchaga

## Abstract

Neuro-inflammation signaling has been identified as an important hallmark of Alzheimer’s disease (AD) in addition to amyloid β plaques (Aβ) and neurofibrillary tangles (NFTs). However, our knowledge of neuro-inflammation is very limited; and the core signaling pathways associated with neuro-inflammation are missing. From a novel perspective, i.e., investigating weakly activated molecular signals (rather than the strongly activated molecular signals), in this study, we uncovered the core neuro-inflammation signaling pathways in AD. Our novel hypothesis is that weakly activated neuro-inflammation signaling pathways can cause neuro-degeneration in a chronic process; whereas, strongly activated neuro-inflammation often cause acute disease progression like in COVID-19. Using the two large-scale genomics datasets, i.e., Mayo Clinic (77 control and 81 AD samples) and RosMap (97 control and 260 AD samples), our analysis identified 7 categories of signaling pathways implicated on AD and related to virus infection: immune response, x-core signaling, apoptosis, lipid dysfunctional, biosynthesis and metabolism, and mineral absorption signaling pathways. More interestingly, most of genes in the virus infection, immune response and x-core signaling pathways, are associated with inflammation molecular functions. Specifically, the x-core signaling pathways were defined as a group of 9 signaling proteins: MAPK, Rap1, NF-kappa B, HIF-1, PI3K-Akt, Wnt, TGF-beta, Hippo and TNF, which indicated the core neuro-inflammation signaling pathways responding to the low-level and weakly activated inflammation and hypoxia, and leading to the chronic neuro-degeneration. The core neuro-inflammation signaling pathways can be used as novel therapeutic targets for effective AD treatment and prevention.

## Introduction

A major challenge limiting effective treatments for Alzheimer’s disease (AD) is the complexity of AD. More than 42 genes/loci have been associated with AD^1,2^. Unfortunately, only few of these genes, like *CD33*^3^, *TREM2*^4^, *MS4A*^5^, are being evaluated as therapeutic targets for AD management^1^. Over 240 drugs have been tested in AD clinical trials, but no new drugs have been approved for AD since 2003^6,7^. One major challenge is that the complicated pathogenesis and core signaling pathways of AD remains unclear. Therefore, it is significant to uncover the core signaling pathways implicated on AD pathogenesis and novel therapeutic targets of AD for identifying effective drugs and synergistic drug combinations (targeting multiple essential targets on the cores signaling network) for AD prevention or treatment.

Our knowledge of the molecular mechanisms and signaling pathways that ultimately lead to the chronic neurodegeneration in AD is limited. For example, there are only a few strong genetic biomarkers for AD that have been identified, including the *APOE, APP, PSEN1/2* genes. However, the signaling consequence of these biomarkers as they relate to the accumulation of dysfunctional A-beta and p-Tau proteins, as well as neuron death and immune response remain unclear. Over the last 10 years, neuro-inflammation and immune signaling have been being identified as the third core feature or a central pathogenesis mechanism of AD^8,9,10,11,12^, in addition to amyloid β plaques (Aβ) and neurofibrillary tangles (NFTs) pathologies. However, our knowledge of neuro-inflammation and immune signaling and their roles in neuro-degeneration is limited, though a set of inflammation and immune genes, like TNF, IL-1beta, IL-6, NFkB have been reported. No computational network analysis has been specifically designed and conducted to uncover and understand the neuro-inflammation and immune signaling pathways systematically. Therefore, it is important to continue to pursue systematic investigations, including the use of network analysis techniques, in order to uncover and understand the details of core signaling pathways, and the core neuroinflammation and immune signaling pathways that are associated the neurodegeneration of AD.

In response to the preceding gap in knowledge, we have systematically sought to identify the potential core signaling pathways causing neuron death and/or degeneration in AD by analyzing the RNA-seq data of human AD samples^13,14^. Instead of identifying the strongly activated molecular signals in the computational network analysis^15^, our unique contribution via this study is to identify the ‘weakly’ activated signaling pathways that may lead to neuron death in a chronic manner. The rationale of focusing on the weakly activated signaling pathways is that the only weakly activated signaling can cause the neuron death/degeneration in a chronic process. Whereas, strongly activated signaling pathways often cause acute disease progression, such as what is observed in a variety of cancers^16,17^ and COVID-19^18,19^. Specifically, we employed the RNA-seq data of neuropathology-free controls and AD samples from two datasets: ROSMAP^13,14^ and Mayo Clinic^20^. Leveraging this data, we then identified all of the weakly activated and inhibited genes with very low fold change thresholds. Subsequently, we conducted network enrichment analyses to identify relevant core signaling pathways. Further, a network inference analysis was conducted to uncover the potential signaling cascades causing neuron death from the activated signaling pathways.

## Results

### Normal and AD tissue samples are barely separable in the gene expression data space

There were 77 normal control subjects and 81 AD cases in Mayo dataset; and 260 normal control samples and 97 AD cases in ROSMAP dataset. The transcripts per million (TPM) values of 16,132 protein coding genes were obtained by applying the Salmon quantification tool^21^ in alignment-based mode using the STAR aligned RNA-seq data. A multidimensional scaling (MDS) model was used to generate the 2D clustering plots of normal control and AD samples in the Mayo and ROSMAP datasets respectively (see **Fig. 1**). As is seen in these visualizations, the normal and AD samples are barely separable, especially in the ROSMAP dataset, which of note, has more normal samples than Mayo dataset.

**Figure 1:**
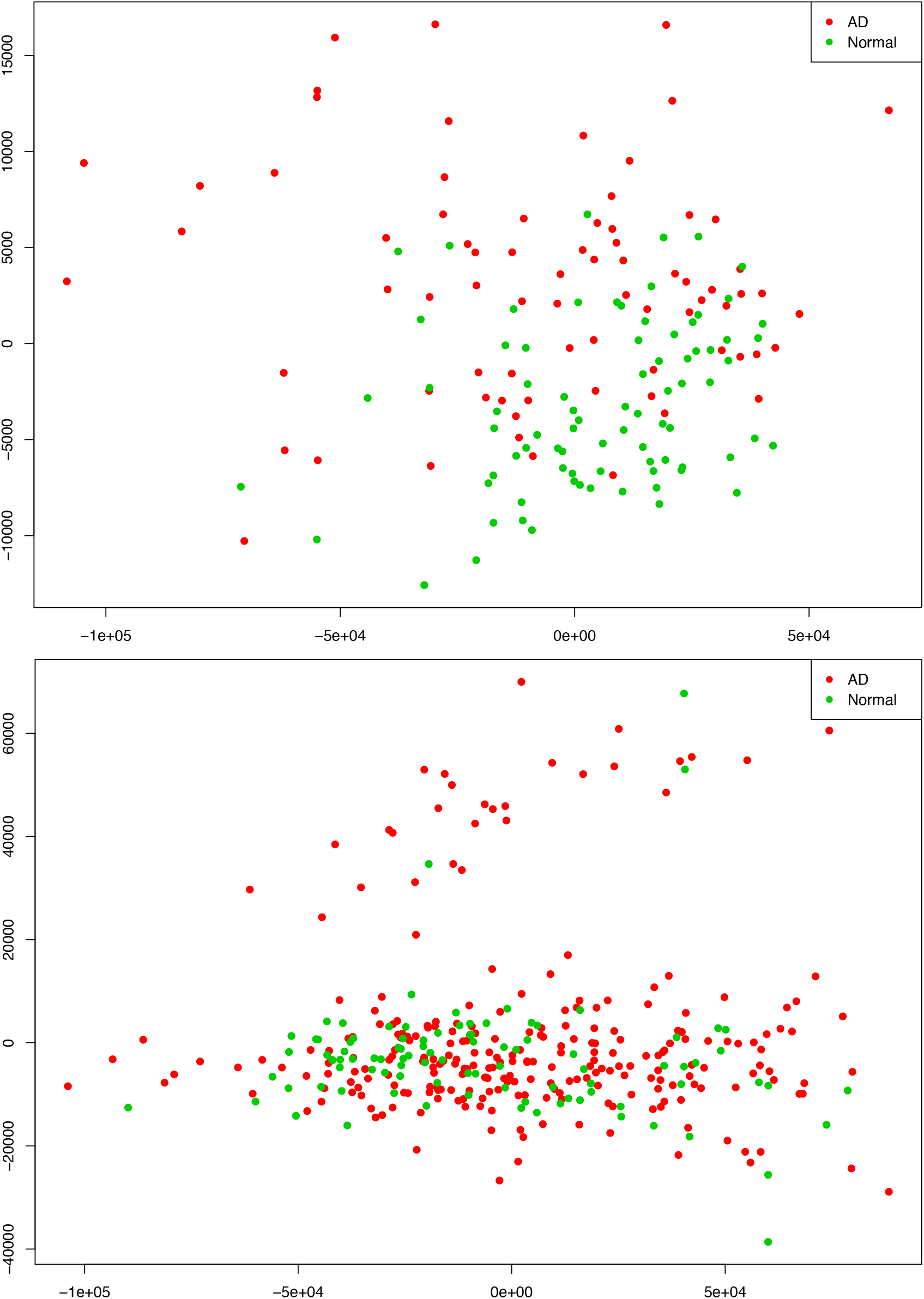
AD and normal control tissue samples are not well separable using an MDS plot on the RNA-seq protein-coding genes in Mayo (**top-panel**) and ROSMAP (**bottom-panel**) datasets.

We further conducted a widely used differential expression analysis method to identify differentially expressed genes (DEGs) between the AD and normal control samples. To identify the common set of DEGs between the two datasets, we applied a number of fold-change and p-value thresholds. As seen in **Table 1** and expected from **Fig. 1**, only a few common up- and down-regulated DEGs were identified with fold change thresholds >= 1.5 and p-value <= 0.05. Even with the fold change threshold >= 1.25, only about 230 up- and about 60 down-regulated genes were identified (out of the 16,132 protein coding genes, ∼1.85%), in both studies. When relaxing both thresholds to fold change >= 1.1 and p-value <= 0.1, 1,120 up-regulated genes and 689 down-regulated genes were identified (∼11.2% of the 161,32 protein-coding genes). Based upon these observations, we hypothesized that the AD-associated signaling pathways are weakly activated or inhibited.

**Table 1:** Epidemiology information of Mayo and RosMap datasets.

| **Mayo** | **Control** | **AD** |  | **RosMap** | **Control** | **AD** |
| --- | --- | --- | --- | --- | --- | --- |
| In Total | 77 | 81 |  | In Total | 97 | 260 |
| Male | 40 | 33 |  | Male | 44 | 82 |
| Female | 37 | 48 |  | Female | 53 | 178 |
| Age Mean (SD) | 82.65 (8.70) | 82.57 (7.62) |  | Age Mean (SD) | 84.24 (6.82) | 90.34 (5.75) |
| APOE_22 | 0 | 0 |  | APOE_22 | 2 | 0 |
| APOE_23 | 12 | 4 |  | APOE_23 | 13 | 22 |
| APOE_33 | 56 | 34 |  | APOE_33 | 72 | 141 |
| APOE_24 | 1 | 0 |  | APOE_24 | 1 | 10 |
| APOE_34 | 8 | 36 |  | APOE_34 | 8 | 83 |
| APOE_44 | 0 | 7 |  | APOE_44 | 1 | 3 |

**Table 1:**
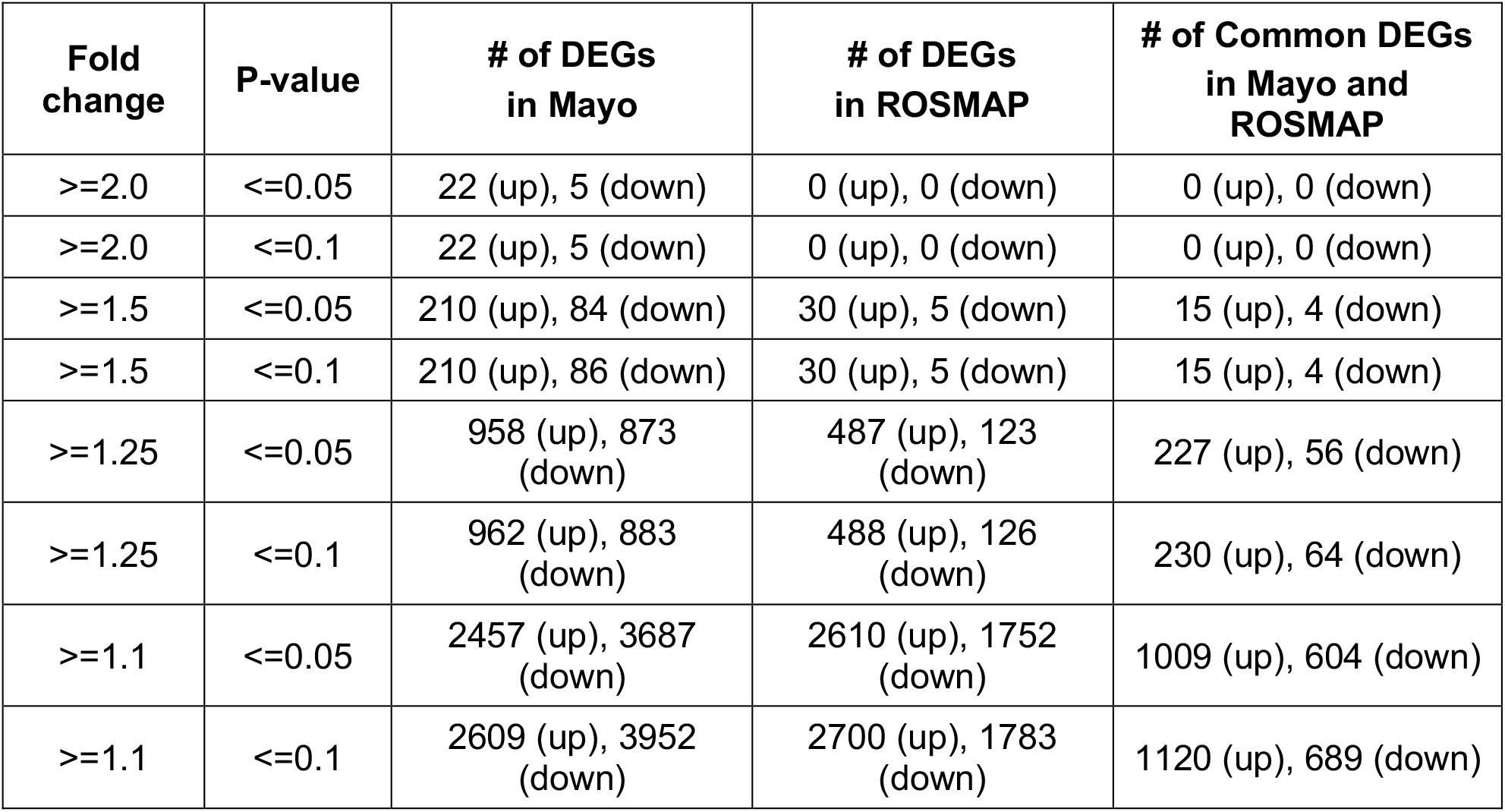
Differentially expressed genes (DEGs) out of 16,132 common protein coding genes between AD and control samples in Mayo and ROSMAP datasets.

| **Fold change** | **P-value** | **# of DEGs in Mayo** | **# of DEGs in ROSMAP** | **# of Common DEGs in Mayo and ROSMAP** |
| --- | --- | --- | --- | --- |
| $\geq 2.0$ | $\leq 0.05$ | 22 (up), 5 (down) | 0 (up), 0 (down) | 0 (up), 0 (down) |
| $\geq 2.0$ | $\leq 0.1$ | 22 (up), 5 (down) | 0 (up), 0 (down) | 0 (up), 0 (down) |
| $\geq 1.5$ | $\leq 0.05$ | 210 (up), 84 (down) | 30 (up), 5 (down) | 15 (up), 4 (down) |
| $\geq 1.5$ | $\leq 0.1$ | 210 (up), 86 (down) | 30 (up), 5 (down) | 15 (up), 4 (down) |
| $\geq 1.25$ | $\leq 0.05$ | 958 (up), 873 (down) | 487 (up), 123 (down) | 227 (up), 56 (down) |
| $\geq 1.25$ | $\leq 0.1$ | 962 (up), 883 (down) | 488 (up), 126 (down) | 230 (up), 64 (down) |
| $\geq 1.1$ | $\leq 0.05$ | 2457 (up), 3687 (down) | 2610 (up), 1752 (down) | 1009 (up), 604 (down) |
| $\geq 1.1$ | $\leq 0.1$ | 2609 (up), 3952 (down) | 2700 (up), 1783 (down) | 1120 (up), 689 (down) |

### Weak inflammation and hypoxia are the potential major factors in the AD brain microenvironment causing neuron cell death

As was noted previously, we believe it is important to identify AD-associated weakly activated signaling pathways, and understand their roles in AD disease progression, as well as their potential roles as targeted for AD therapeutics. Among the 1,120 common up-regulated genes (identified from Mayo and ROSMAP datasets), 417 genes were included in the 311 KEGG signaling pathways. To this end, we first conducted an enrichment analysis of KEGG signaling pathways using Fisher’s exact test applied to the 417 up-regulated genes. **Table 2** showed the enriched signaling pathways with p-value <= 0.15. We then clustered these activated signaling pathway empirically into 7 categories (see **Fig. 2**). Using these 417 up-regulated genes, the first principal component values in the MDS analysis of the AD and control samples were used to compared the difference in AD and control samples. The OR, absolute beta values and p-values of logistic regression analysis (see **Table 2**) indicated that these selected genes (p-value=1.22×10^−13^ (Mayo) and p-value=4.2×10^−6^ (ROSMAP)) can separate the AD and control samples much better than using all protein genes (p-value=0.036(Mayo) and p-value=0.027 (ROSMAP)) in the two datasets respectively. The bar-plots were also provided in **Fig. 2**, which indicated that the control and AD samples are more separable using the selected genes.

**Table 2:** Odds ratio (OR), beta and p-values of logistic regression using all gene and 417 up-regulated genes.

|  | All genes |  |  | 417 genes |  |  |
| --- | --- | --- | --- | --- | --- | --- |
|  | OR | abs(beta) | p-value | OR | abs(beta) | p-value |
| **Mayo** | 1.42 | 0.35 | 0.037 | 5.9 | 1.78 | $9.5 \times 10^{-9}$ |
| **RosMap** | 1.31 | 0.27 | 0.027 | 1.9 | 0.63 | $9.7 \times 10^{-5}$ |

**Figure 2:**
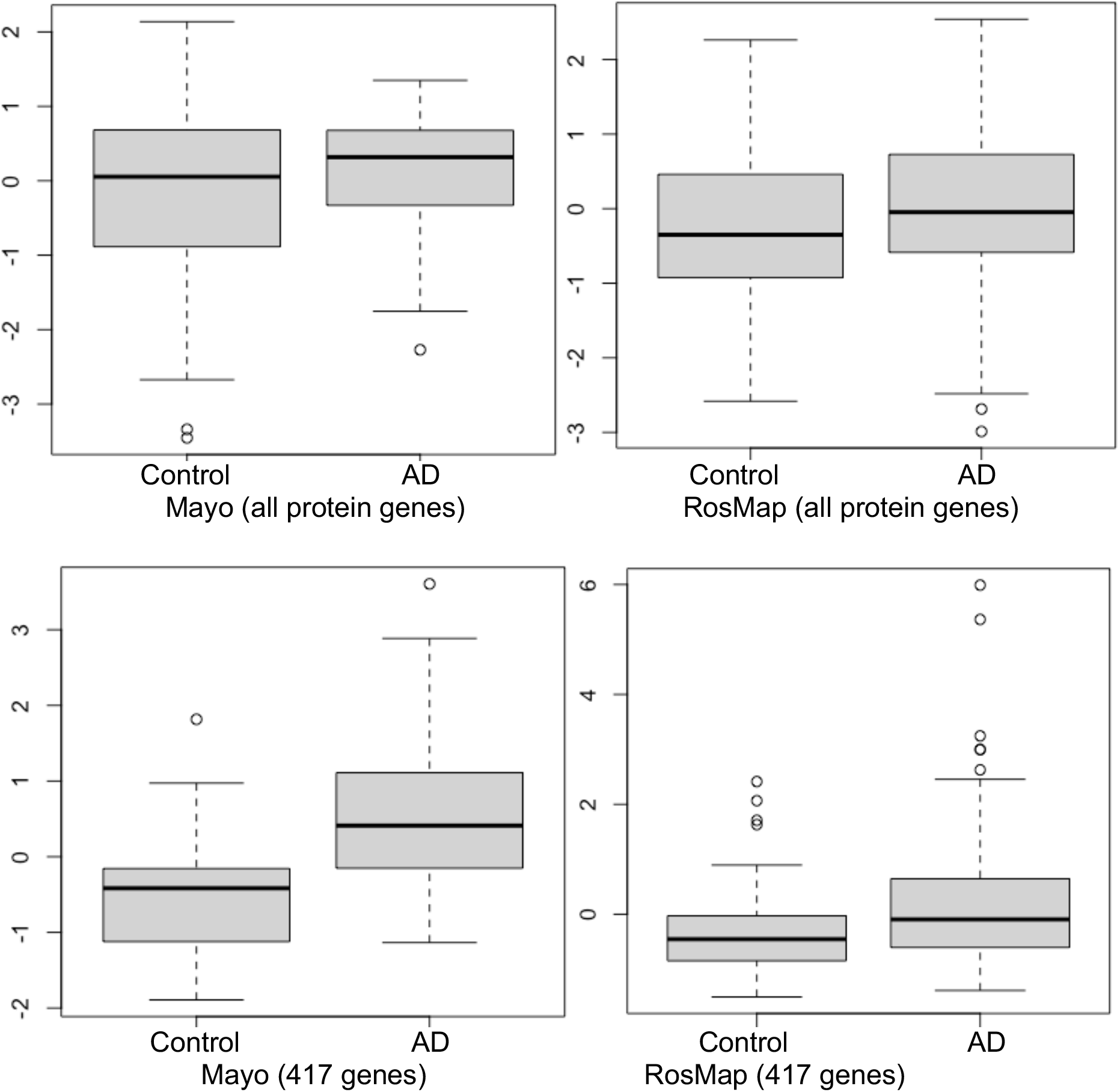
Box-plots of the first principal component of MDS analysis in control and AD cases. **Left and right columns** are Mayo and RosMap samples respectively. **Upper and lower panels** represent the MDS analysis using all genes and 417 up-regulated genes respectively.

As seen in our results (see **Fig.3** and **Table 3**), a set of signaling pathways were activated, such as those involved in virus infection signaling (including: Epstein-Barr virus, Human T-cell leukemia virus 1 infection, Legionellosis, Pathogenic Escherichia coli infection, Staphylococcus aureus infection, Yersinia infection, Human cytomegalovirus infection, Human papillomavirus infection, Malaria, Human immunodeficiency virus 1 infection, Rheumatoid arthritis, and Inflammatory bowel disease [IBD]). There are 111 genes (out of the 417 up-regulated genes) in common across these pathways highlighting a set of core genes implicated on these processes. These results indicated that weakly activated inflammation related signaling pathways, like inflammation, cytokine, and immune response, may be represent activated signaling pathways in the AD brain microenvironment.

**Figure 3:**
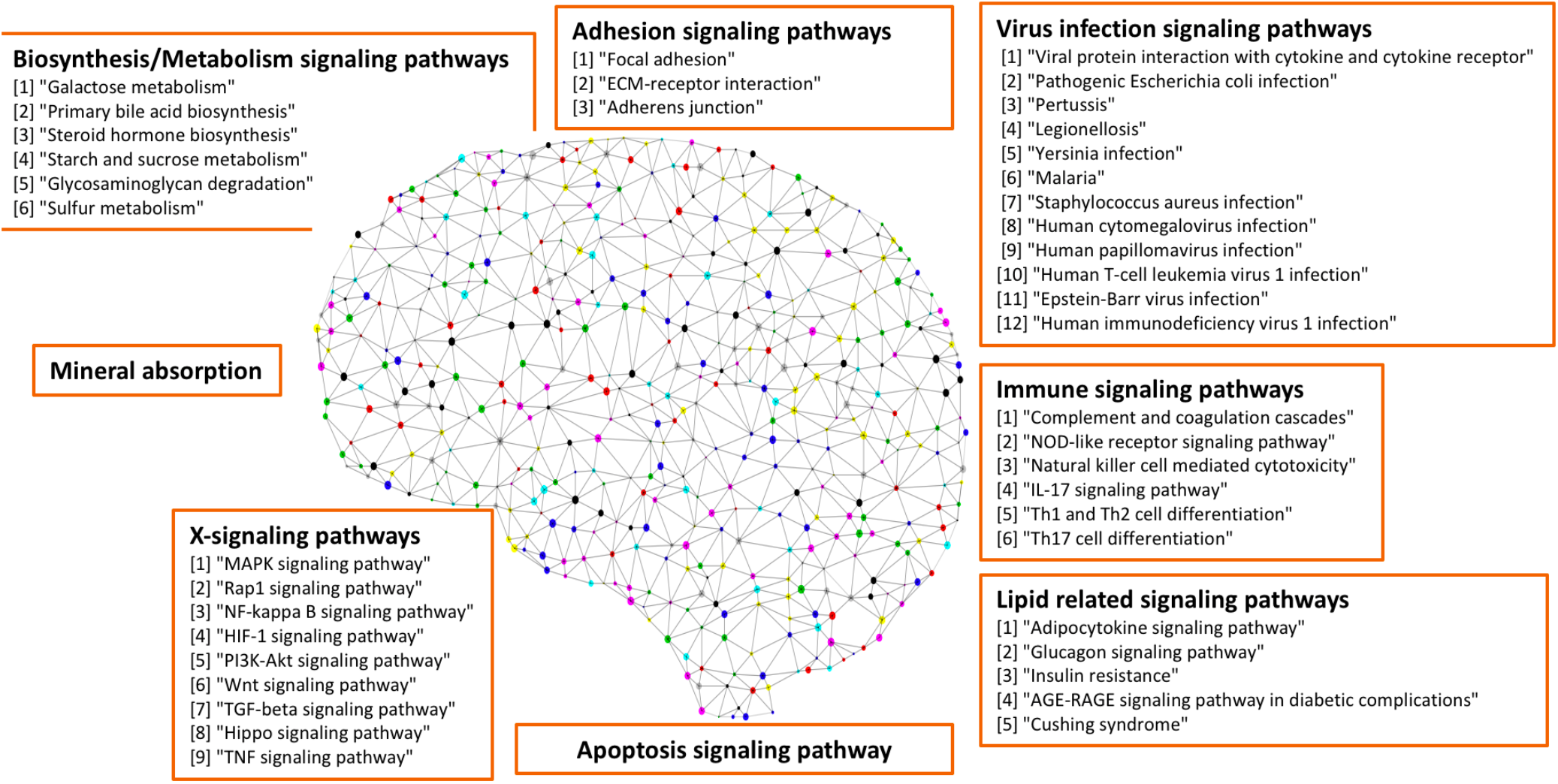
Seven categories of weakly activated signaling pathways in AD.

**Table 3:**
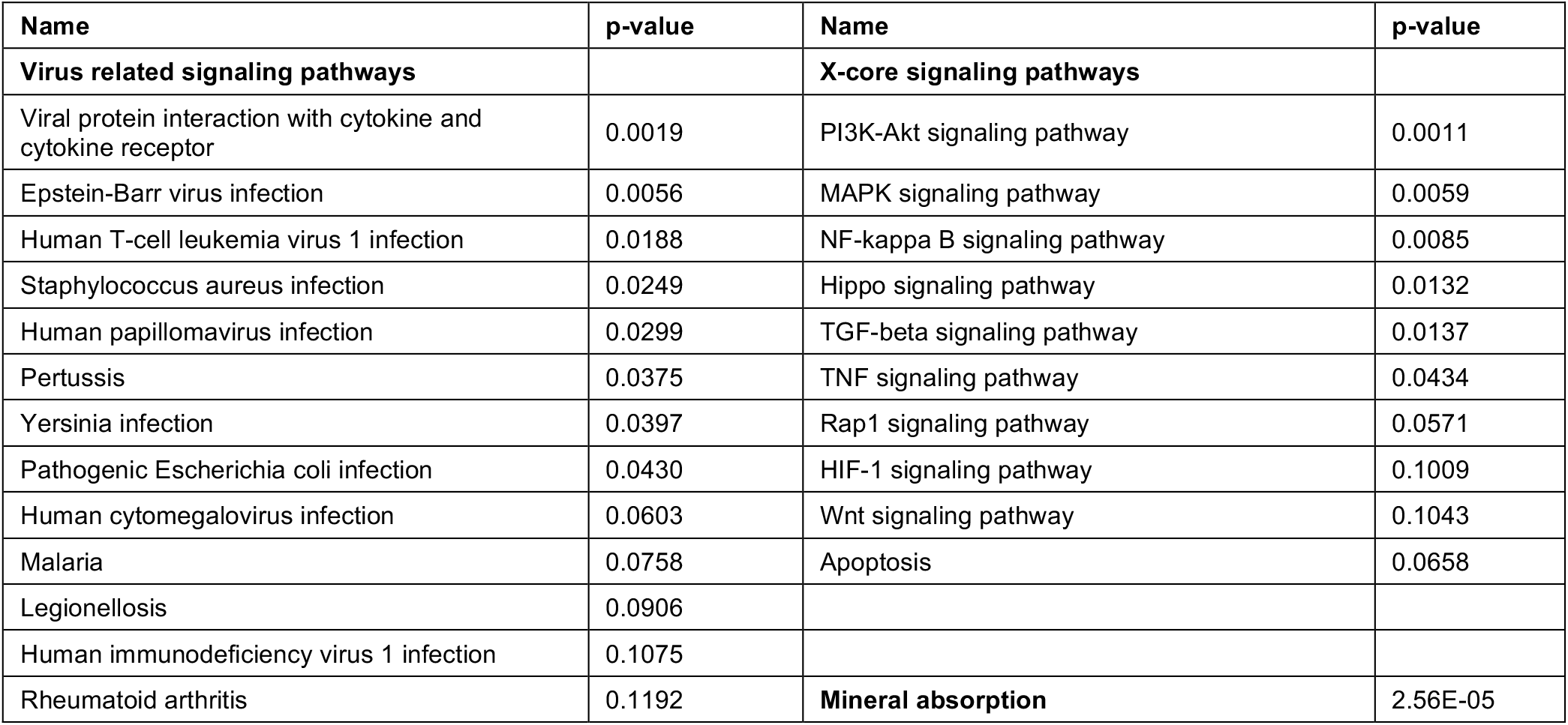
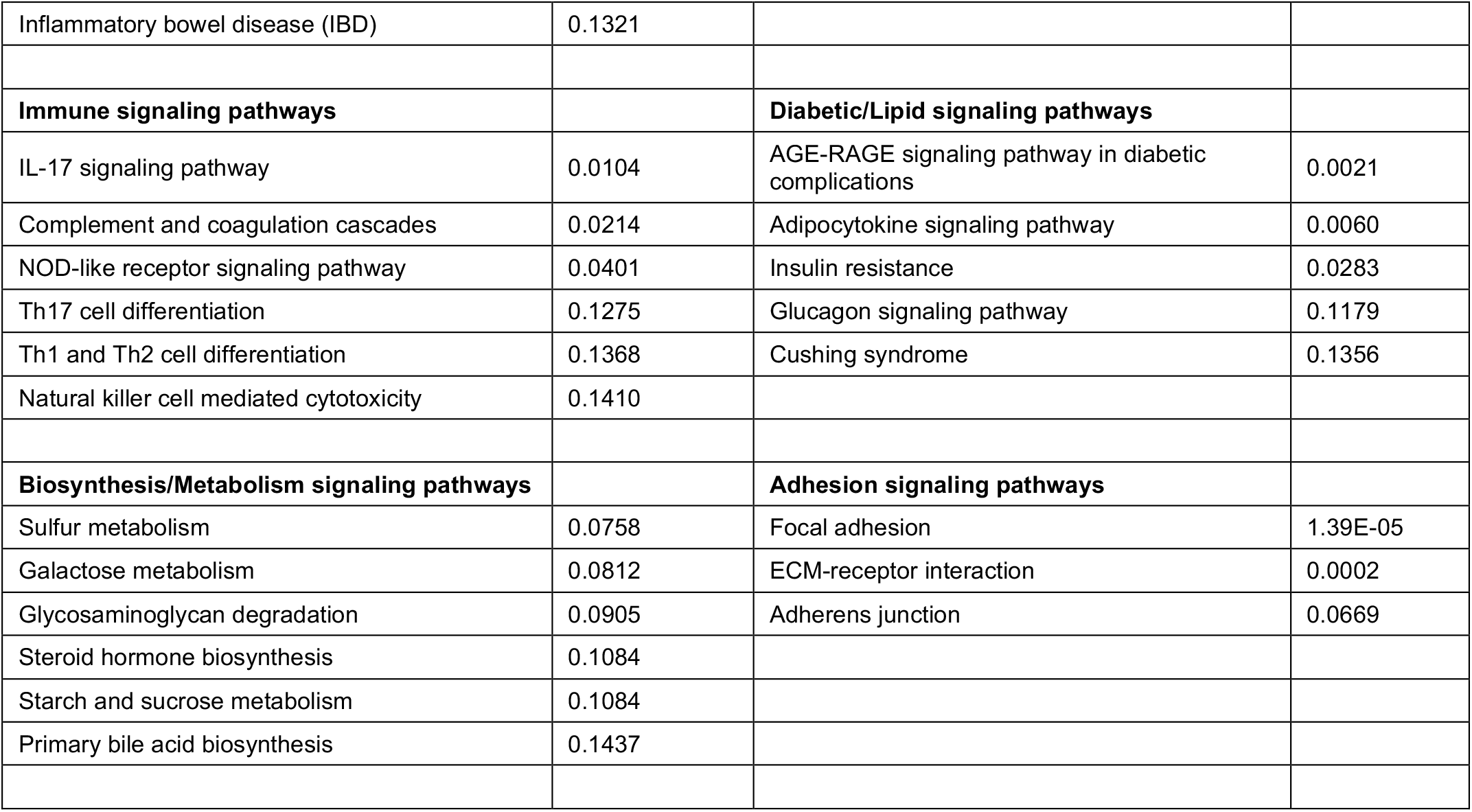
The seven categories of enriched KEGG signaling pathways.

In addition, a group of activated signaling pathways or factors that are not clustering to a specific biological function or disease (referred to as the x-signaling pathway: the Hippo, PI3K-Akt, AGE-RAGE, MAPK, Adipocytokine, NF-kappa B, IL-17, TGF-beta, NOD-like receptor, TNF, Apoptosis, HIF-1 and Wnt signaling pathways, as well as apoptosis signaling) were identified. **Fig. 4** shows the associations between these up-regulated genes and activated signaling pathways. As seen in **Fig. 4**, a set of genes in the center areas of the network are associated with a set of signaling pathways, which could represent therapeutic signaling targets that could be used to inhibit or otherwise perturb these activated signaling pathways. In addition, there are a number of metabolisms signaling pathways, like Sulfur metabolism, Galactose metabolism, Starch and sucrose metabolism, Steroid hormone biosynthesis, Glycosaminoglycan degradation, implicated in this model. Moreover, Th1/2/17 (T helper, CD4+ cells) cell differentiation signaling was activated. Similarly, the natural killer cell mediated cytotoxicity signaling pathways were also activated. **Table S1** lists these associated up-regulated genes and the involved signaling pathways. All the observations suggest a potential novel hypothesis that the external inflammation, immune signaling and hypoxia signaling in AD microenvironment activated the *MAPK, PI3K-Akt and mTOR* signaling pathways, and then activated the *HIF-1* signaling pathway. However, the activation of *HIF-1* may fail to bring enough oxygen to protect against hypoxic injury to the involved neurons. The dysfunction of blood vessel functions, leading to hypoxia, might be partially indicated by the recent study showing that blood and cerebrospinal fluid flow cleaning the brain during sleeping^22^.

**Figure 4:**
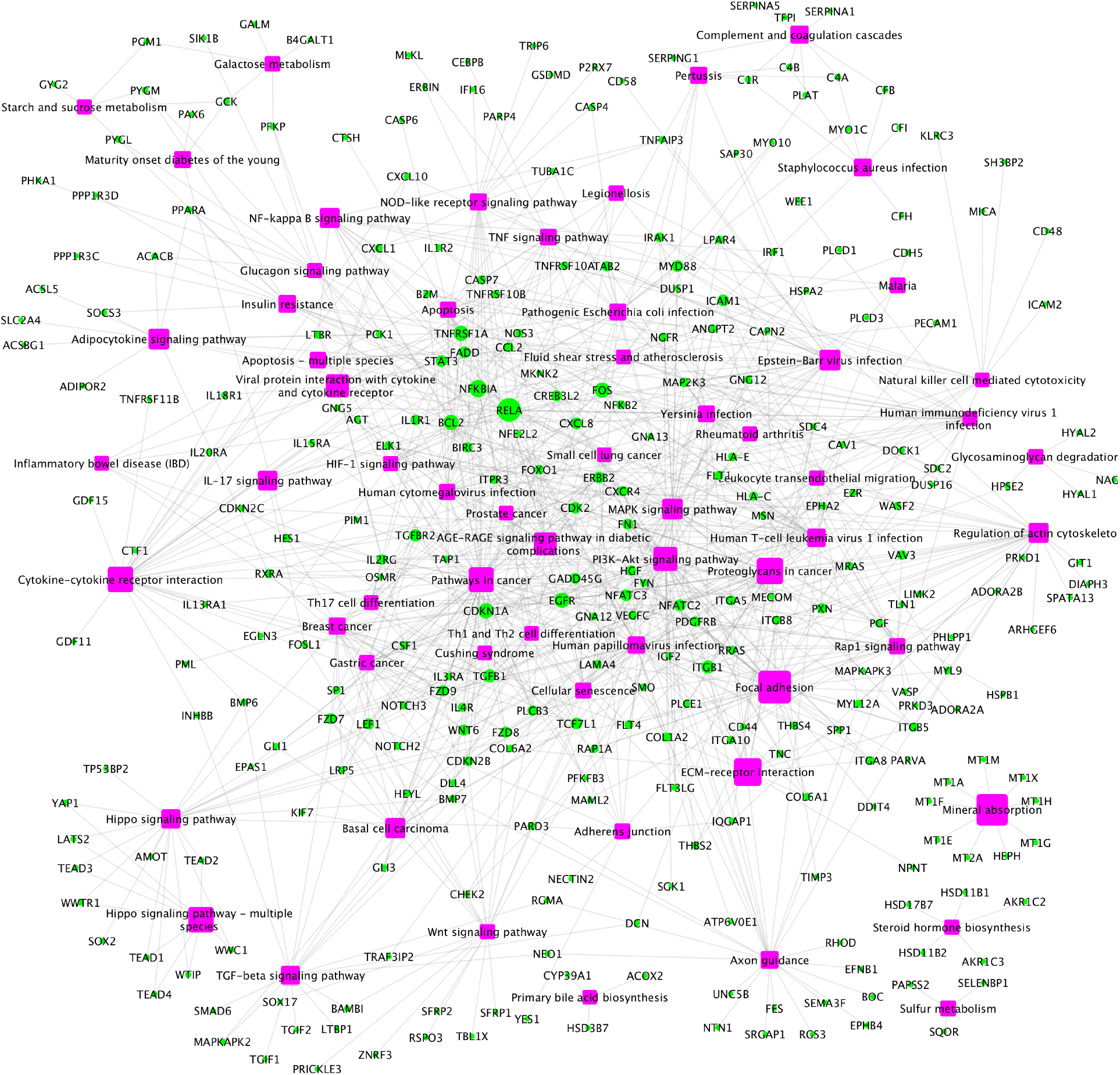
The up-regulated gene-pathway interaction network, including 1021 interactions between 291 up-regulated genes and 61 enriched pathways.

### Weak inflammation and hypoxia are the major factors in the AD brain microenvironment causing neuron cell death

As was introduced above, many genes that are activated as a function of virus infection, immune response and the x-core signaling pathways are inflammation related genes. It is well known that virus infection and immune response signaling pathways respond to inflammation. Our analyses identified 1043 inflammation response genes in the gene ontology (GO) database (GO:0006954), that includes 492 genes in the KEGG signaling pathways. Interestingly, among the 417 up-regulated genes, 66 genes were inflammation related. The p-value of observing the 66 up-regulated inflammation signaling targets from 417 up-regulated genes identified in the AD vs control samples was 8.34E-05 (calculated using the fishers’ exact test), and the odds ratio (OR) = 1.77, which indicate that the activation of inflammation signaling is concomitant with AD progression. Furthermore, there are 66 overlapping up-regulated genes spanning the virus infection (from 111 up-regulated genes) and x-core signaling pathways (from 136 up-regulated genes), which indicate that the x-core signaling pathways are the likely pathways being activated in response to this inflammation. In addition, the activation of HIF-1 signaling pathway indicates the presence of hypoxia in the AD brain environment.

To further investigate the network signaling cascades involving inflammation and apoptosis genes, we conducted the network analysis incorporating the activated signaling pathways and apoptosis signaling genes. As seen in **Fig. 5**, the potential signaling cascades linking the up-regulated inflammation related genes in virus infection and X-core signaling pathways to the activated apoptosis signaling targets. Among the 338 signaling network genes in **Fig. 5**, there are 18 reported GWAS genes (with p-value <= 1.0×10^−5^): *PIK3CB, AKT3, RAF1, MAPK10, PPP2R2B, ERBB4, MECOM, IL1R1, MYD88, CAMK2D, GNB4, VAV3, PRKD3, PRKCE, THRB, FN1, LTBP1 WWTR1*, which were reported in the GWAS analysis ^26^. Further, we also compared the distance distribution among the inflammation related up-regulated genes and apoptosis genes as shown in **Fig. 6**. As can be seen, the inflammation signaling genes are much closer, based on the shortest path metric calculated using the Dijkstra’s algorithm, on the signaling network, (see green, blue and red nodes) to the apoptosis genes compared with other signaling genes (see gray lines). These results indicate a potential signaling interactions between the inflammation signaling genes and apoptosis signaling. In other words, the results suggest a potential association that the weak inflammation and hypoxia signaling in the AD brain environment led to chronic neurodegeneration process via the activation of the x-core signaling pathways. Therefore, drugs and drug combinations that can perturb the X-core signaling pathways have the potential to be effective for AD prevention and treatment.

**Figure 5:**
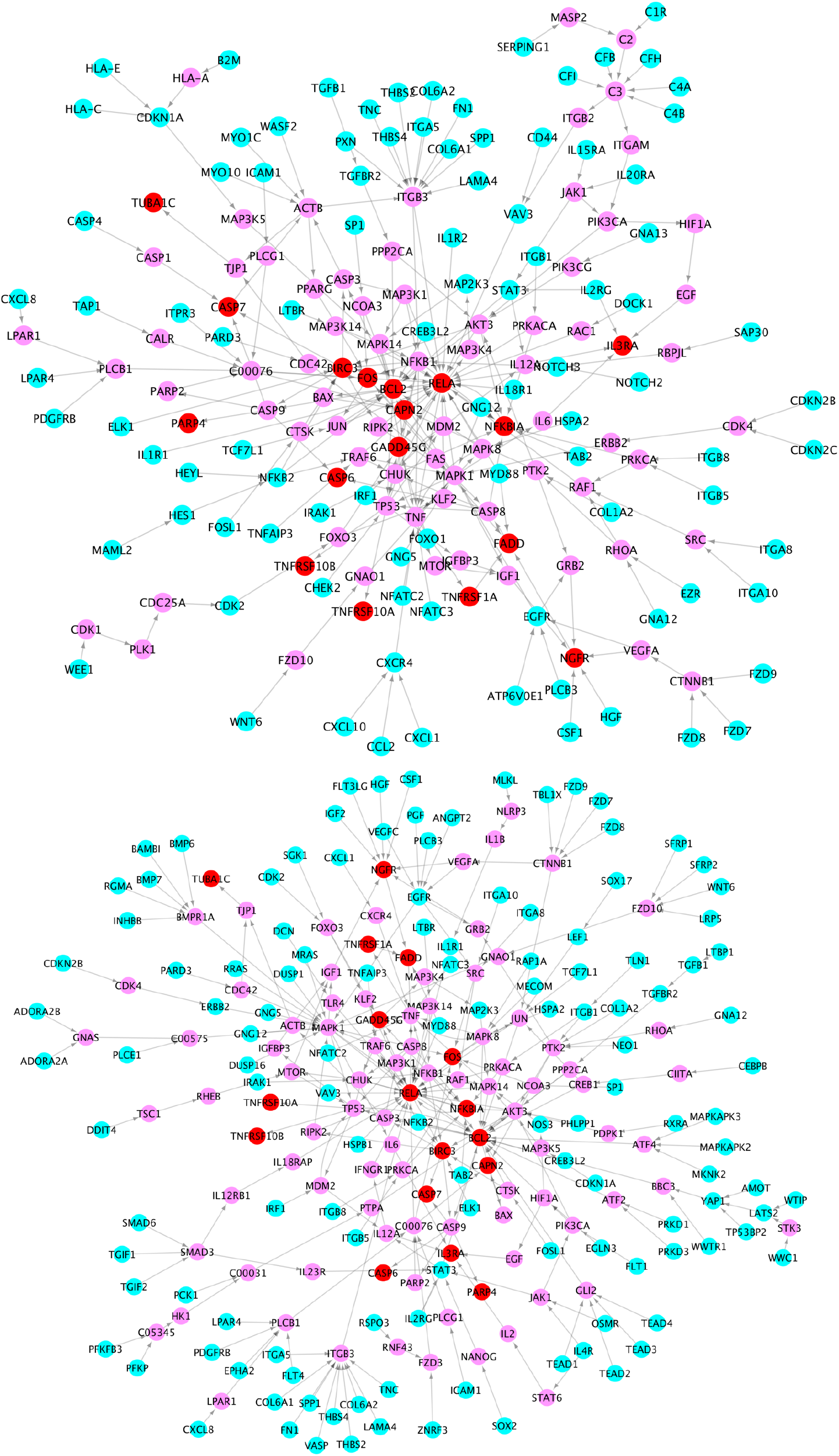
Signaling cascades linking the up-regulated signaling genes in the virus infection pathways (cyan) (**top**) and x-core signaling pathways (**bottom**) to the up-regulated apoptosis signaling genes (red) via the linking genes (pink).

**Figure 6:**
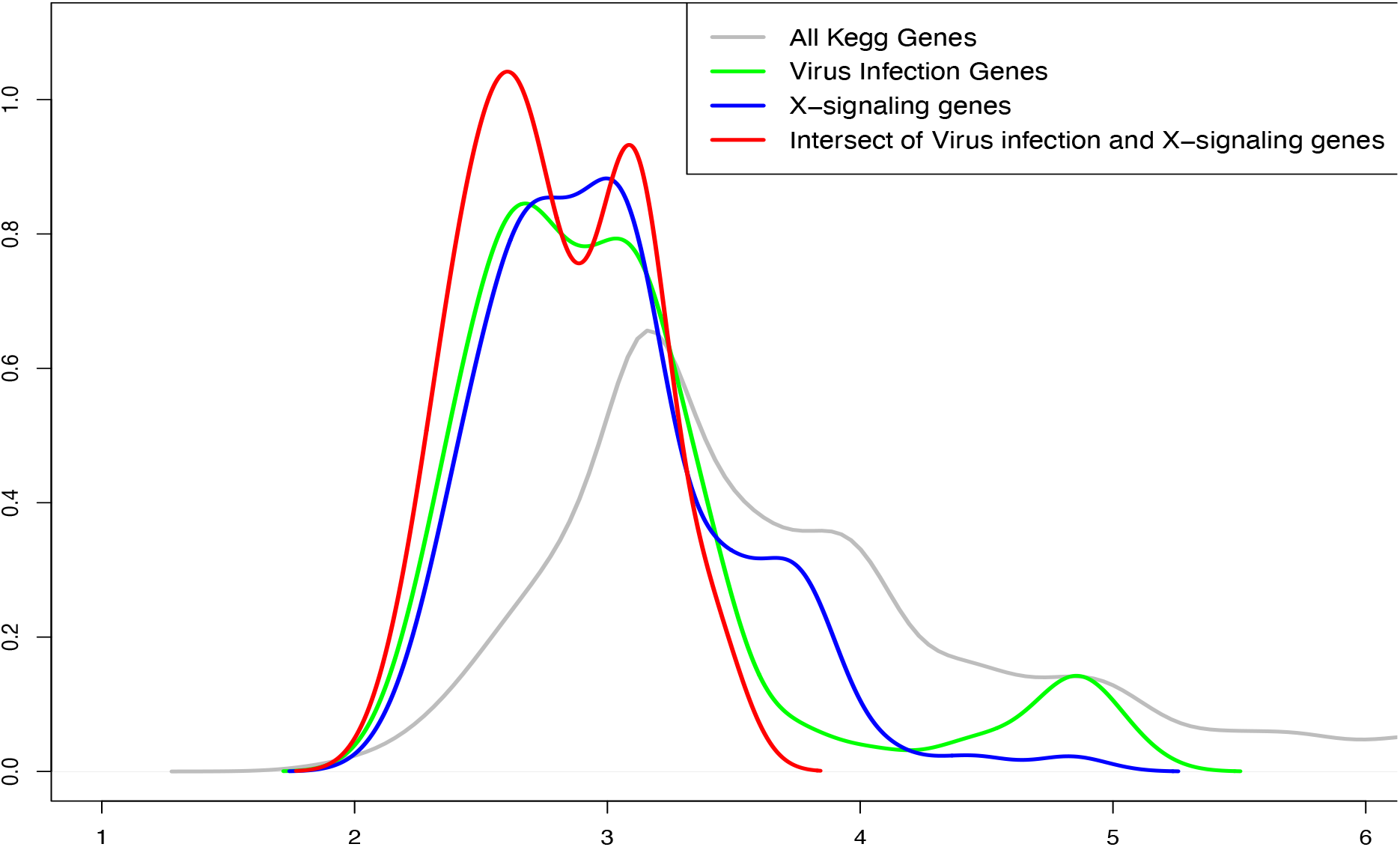
The up-regulated genes in the inflammation related signaling pathways, including virus infection, immune response and x-core signaling pathways. As seen, the inflammation signaling genes are much closer (see green, blue and red) to the apoptosis genes compared with other signaling genes (see gray lines).

### Activated TNF signaling might lead the programmed apoptosis of neurons

Of note, our results show that among the X-core signaling pathways, the TNF signaling pathways are also activated. Particularly, the TNF (Tumor Necrosis Factor) receptors (TNFRSF1A TNFRSF10A, and TNFRSF10B) were up-regulated (see **Table 4**). We reconstructed these signaling pathway linking the TNF receptors to the up-regulated genes in TNF and apoptosis signaling pathways (see in **Fig. 7**). As seen, the activation of these TNF signaling pathway might be one possible molecular mechanism causing the activation of apoptosis signaling via the CASP6, CASP7 cascades.

**Table 4:** up-regulated genes in TNF and apoptosis signaling pathways.

|  |  |
| --- | --- |
| Apoptosis | BCL2, RELA, BIRC3, FADD, GADD45G, TNFRSF1A, NFKBIA, TNFRSF10B, CAPN2, TUBA1C, IL3RA, CTSH, FOS, CASP6, CASP7, TNFRSF10A, PARP4 |
| TNF signaling pathway | RELA, BIRC3, FADD, MAP2K3, TNFRSF1A, NFKBIA, CREB3L2, FOS, CASP7, MLKL, IRF1, CEBPB |

**Figure 7:**
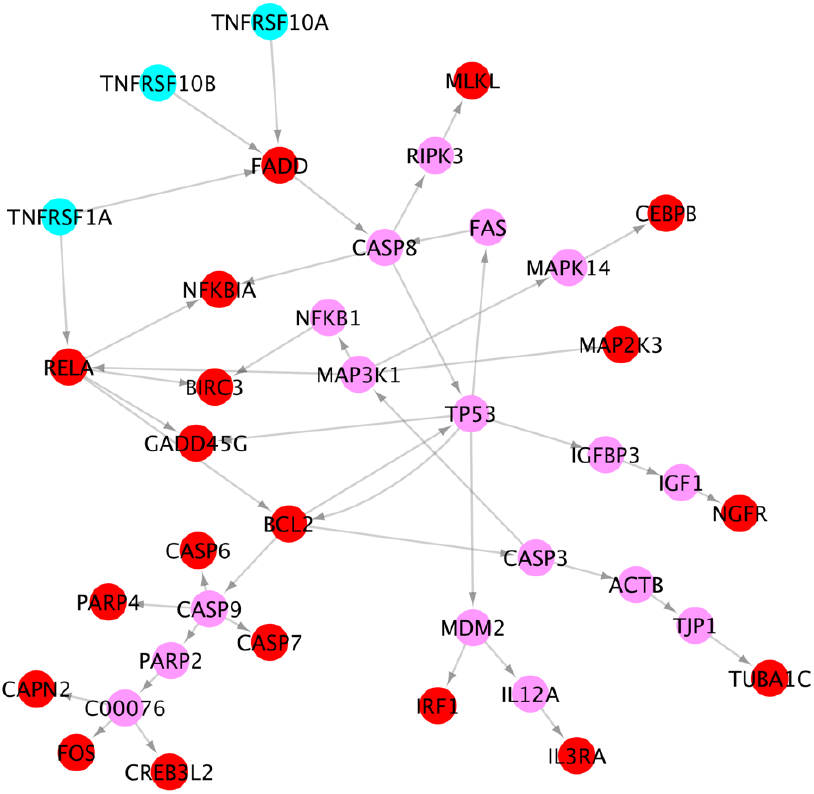
Signaling cascades, causing neuron death, from the 3 TNF receptors (cyan) to the up-regulated genes (red) in TNF and apoptosis signaling pathways via the linking genes (pink).

### Circadian entrainment, addiction, Neuroactive Neuroactive ligand-receptor, Synaptic vesicle cycle, Fat acid biosynthesis, mTOR and oxidative phosphorylation signaling pathways were down-regulated inhibited in AD tissues

We also conducted pathway enrichment analyses using 143 down-regulated genes in KEGG signaling pathways. There are far fewer down-regulated genes and lower inhibition down-regulated of KEGG signaling pathways, compared with up-regulated genes (see **Table 5**). As shown in **Fig. 8**, the circadian entrainment, addition, Neuroactive neuroactive ligand-receptor, Synaptic vesicle cycle, Fat acid biosynthesis and oxidative phosphorylation genes pathways were inhibited in AD, which is associated with the down-regulated Ras and cAMP signaling pathways.

**Table 5:** The 30 down-regulated KEGG signaling pathways.

| Pathway Names | p-value |
| --- | --- |
| **Neuron related signaling pathways** |  |
| Neuroactive ligand-receptor interaction | 4.35E-06 |
| Circadian entrainment | 0.003 |
| Glutamatergic synapse | 0.009 |
| Synaptic vesicle cycle | 0.017 |
| SNARE interactions in vesicular transport | 0.036 |
| Dopaminergic synapse | 0.040 |
| GABAergic synapse | 0.148 |
| **Addiction signaling pathways** |  |
| Nicotine addiction | 0.003 |
| Alcoholism | 0.006 |
| Morphine addiction | 0.075 |
| **Fatty acid signaling pathways** |  |
| Fatty acid elongation | 0.062 |
| Fatty acid biosynthesis | 0.132 |
| **Signaling transduction pathways** |  |
| cAMP signaling pathway | 0.023 |
| mTOR signaling pathway | 0.023 |
| Ras signaling pathway | 0.083 |
| **Infection signaling pathways** |  |
| Oxidative phosphorylation | 4.05E-05 |
| Vibrio cholerae infection | 0.0002 |
| Amphetamine addiction | 0.0009 |
| Epithelial cell signaling in Helicobacter pylori infection | 0.0009 |
| **Metabolism signaling pathways** |  |
| Taurine and hypotaurine metabolism | 0.038 |
| Butanoate metabolism | 0.046 |
| Cysteine and methionine metabolism | 0.060 |
| Retinol metabolism | 0.088 |
| **Other signaling pathways** |  |
| Valine, leucine and isoleucine biosynthesis | 0.007 |
| Phagosome | 0.018 |
| Retrograde endocannabinoid signaling | 0.056 |
| Long-term potentiation | 0.071 |
| Salivary secretion | 0.078 |
| Gastric acid secretion | 0.083 |
| Phototransduction | 0.086 |

**Figure 8:**
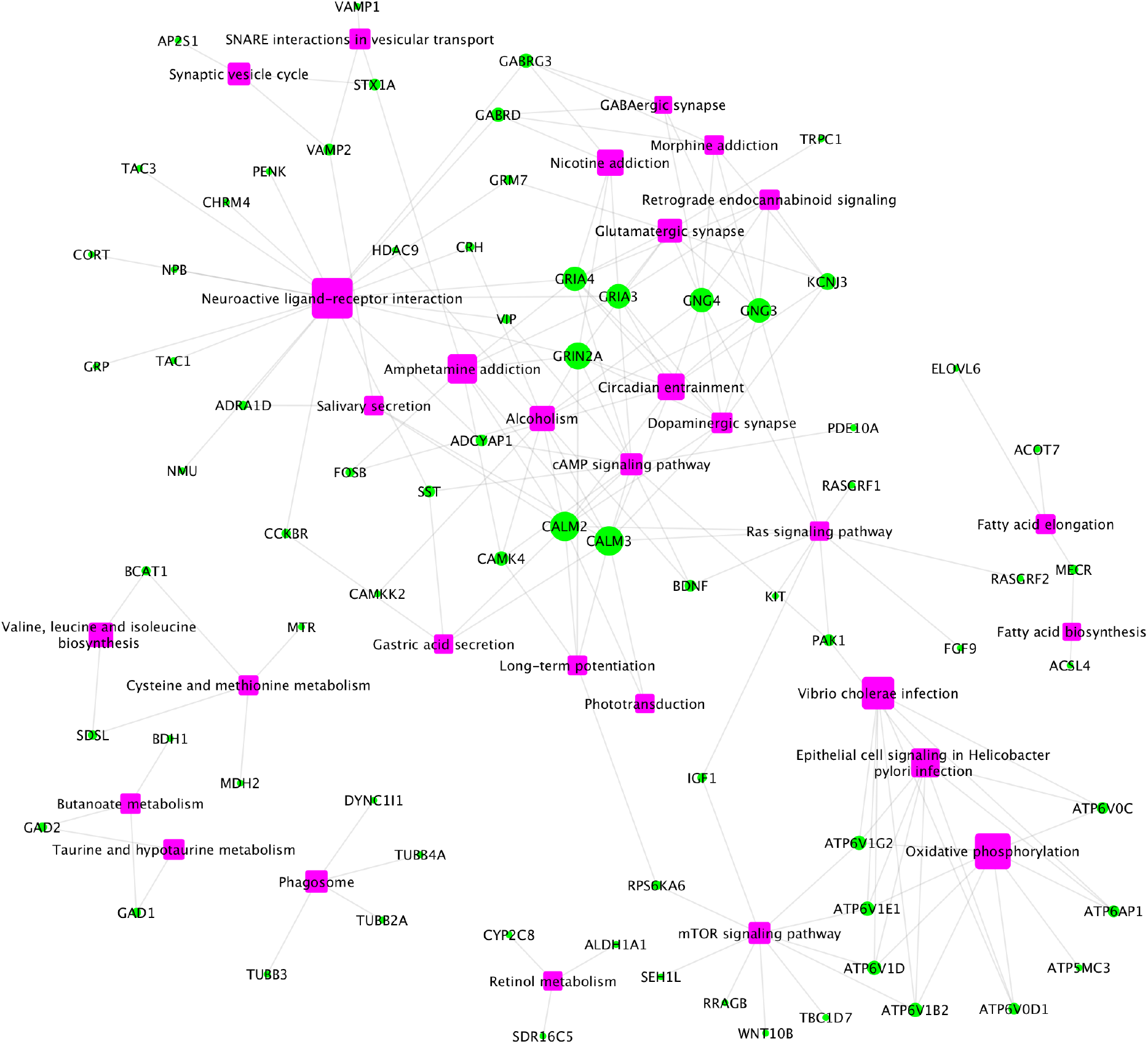
The down-regulated gene-pathway interaction network, including 183 interactions between 73 up-regulated genes and 30 enriched pathways.

The dysfunctional circadian entrainment signaling pathway were reported to be associated with AD, and might be associated with the rhythmic spinal fluid washing over brain during deep sleep. In addition, the mTOR signaling and oxidative phosphorylation signaling pathways were down-regulated. Moreover, fat acid biosynthesis and elongation were also inhibited.

## Methods

### Gene expression data analysis of Apoe4/4 genotype AD samples

In this study, 77 normal tissue samples and 81 AD tissue samples in Mayo dataset; and 97 normal samples and 260 AD samples in ROSMAP dataset were used. Both datasets were processed and aligned separately using reference genome GRCh38 and GENCODE 33 annotation including the ERCC spike-in annotations. We excluded ALT, HLA, and Decoy contigs from the reference genome due to the lack of RNA-Seq tools that allow to handle these regions properly. To obtain gene expression data, all read sequences from both datasets were first mapped to the reference genome using STAR (v.2.7.1a)^23^.Transcripts per million (TPM) values of 16,132 common protein coding genes were then obtained in the two datasets by applying the Salmon quantification tool^21^ in alignment-based mode using the aligned RNA-seq data.

### Differentially expressed genes

To identify the up- and down-regulated genes in AD samples vs normal control samples, the edgeR^24^ tool, using the negative binomial (NB) statistical model, was applied to the TPM values.

### Inflammation genes

A set of inflammation genes were obtained by extracting genes from the inflammatory response category as defined in the Gene Ontology (GO:0006954)^25^. Subsequently, 485 inflammation genes were obtained from the 5,191 KEGG signaling genes.

### AD GWAS data

The GWAS data of AD was obtained from niagads database^26^ (https://www.niagads.org/igap-rv-summary-stats-kunkle-p-value-data). The Stage 1 P-Value Data (updated by February 26, 2019) and Stage 2 P-Value Data (updated by February 27, 2019) were downloaded. The 553 candidate GWAS genes and also available in the KEGG signaling pathways were obtained by a filter with p-value <= 1.0×10^−5^.

### KEGG signaling pathway enrichment analysis

The KEGG signaling pathways consist of 311 signaling pathways^27,28^. There are 59,242 signaling interaction among 5,191 genes in these pathways, which were used for network enrichment analysis and network inference analysis in this study. For the network enrichment analysis, a Fisher’s exact test^29,30^ was used based upon the up-regulated genes.

### KEGG signaling network inference analysis

To infer the signaling cascades among a set of genes of interest, we developed a network inference approach. First, we divided the genes into two groups: signaling sources (like the inflammation signaling genes), and signaling targets (like the apoptosis signaling genes). Second, a signaling network was constructed by linking the signaling source genes to the signaling target genes iteratively. Specifically, the signaling source genes was used as the initial signaling source nodes set: V0. The signaling target genes were used as the target nodes set: V1. In the iterative process, the shortest signaling cascades/paths between the nodes in V0 and V1 were calculated and identified: *P*_*ij*_ *= <g*_*i*_, *g*_*k1*_, *g*_*k2*_, *…, g*_*j*_*>, where g*_*i*_ *belongs to V0, and g*_*j*_ *belongs to V1*. Third, all of the genes on the signaling path *P*_*ij*_ and belong to V1 were selected and added to V0, and removed from V1. This process was repeated until all the genes were added to V0.

## Discussion and conclusion

Neuro-inflammation and immune signaling have been being identified as an important pathogenesis mechanism of AD, in addition to amyloid β plaques (Aβ) and neurofibrillary tangles (NFTs) pathologies. However, our knowledge of neuro-inflammation and immune signaling and their roles in neuro-degeneration is limited, though a set of inflammation and immune genes, like TNF, IL-1beta, IL-6, NFkB have been reported. Recently, the network analysis models were proposed to identify the potential dysfunctional signaling pathways and biomarkers using the related RNA-seq datasets. For example, the molecular signatures and networks under different brain regions were reported using integrative co-expression network analysis, and the myelin signaling dysregulation was identified in AD^31,32^. In addition, the co-splicing network using the WGCNA (co-expression network analysis model) was conducted to identified the altered splicing in AD, which indicated that the altered splicing is the mechanism for the effects of the AD related CLU, PTK2b and PICALM alleles33. Moreover, the molecular subtypes and potential driver genes, like CABRB2, LRP10, ATP6V1A, of AD were identified by combing key driver analysis (KDA) and multiscale embedded gene expression network analysis (MEGENA)^34,35,36^. However, neuro-inflammation and immune signaling pathways have not been systematically uncovered and analyzed in these reported computational models. Compared with these reported studies, our unique contribution is the novel discovery of essential neuro-inflammation and immune signaling genes and signaling interactions using systematic network analysis models, which indicates potentially novel targets and mechanisms of neuro-inflammation and immune signaling in neuro-degeneration.

Specifically, we propose a novel hypothesis that weakly activated neuro-inflammation signaling pathways can cause neuro-degeneration in a chronic process; whereas, strongly activated neuro-inflammation often cause acute disease progression like in COVID-19. Consequently, from a novel perspective, i.e., investigating the weakly activated molecular signals (rather than the strongly activated molecular signals), in this study, we uncovered the core neuro-inflammation signaling pathways in AD. To the best of our knowledge, it is the first time to systematically uncover the core neuro-inflammation signaling pathways based on the transcriptomic data of AD. The neuro-inflammation signaling pathways, including the virus infection, immune response, x-core signaling pathways, apoptosis signaling pathways. indicated that such weak inflammation may lead to the activation of x-core signaling pathways and the ultimate apoptosis of neurons. As a result, we hypothesize that drugs and drug combination inhibiting the neuro-inflammation signaling pathways could be potentially effective for AD prevention and treatment. Moreover, it is interesting to investigate the detailed signaling cascades of the x-core signaling pathways, including the MAPK, Rap1, NF-kappa B, HIF-1, PI3K-Akt, Wnt, TGF-beta, Hippo and TNF signaling pathways. And it is important to study their roles in Aβ plaques and tau tangles as well as neuro-degeneration.

## Acknowledgement

This work is partially supported by National Institute of Ageing (NIA) R56AG065352 to Dr. Li. We thank all the participants and their families, as well as the many involved institutions and their staff. Funding: This work was supported by grants from the National Institutes of Health (R01AG044546 (CC), P01AG003991(CC, JCM), RF1AG053303 (CC), RF1AG058501 (CC), and U01AG058922 (CC), and chuck zuckerberg initiative (CZI). This work was supported by access to equipment made possible by the Hope Center for Neurological Disorders, and the Departments of Neurology and Psychiatry at Washington University School of Medicine. CC receives research support from: Biogen, EISAI, Alector and Parabon. CC is a member of the advisory board of Vivid Genomics, Halia Therapeutics and ADx Healthcare. The remaining authors declare no competing interests.

## Supplementary Tables

**Table ST1:**
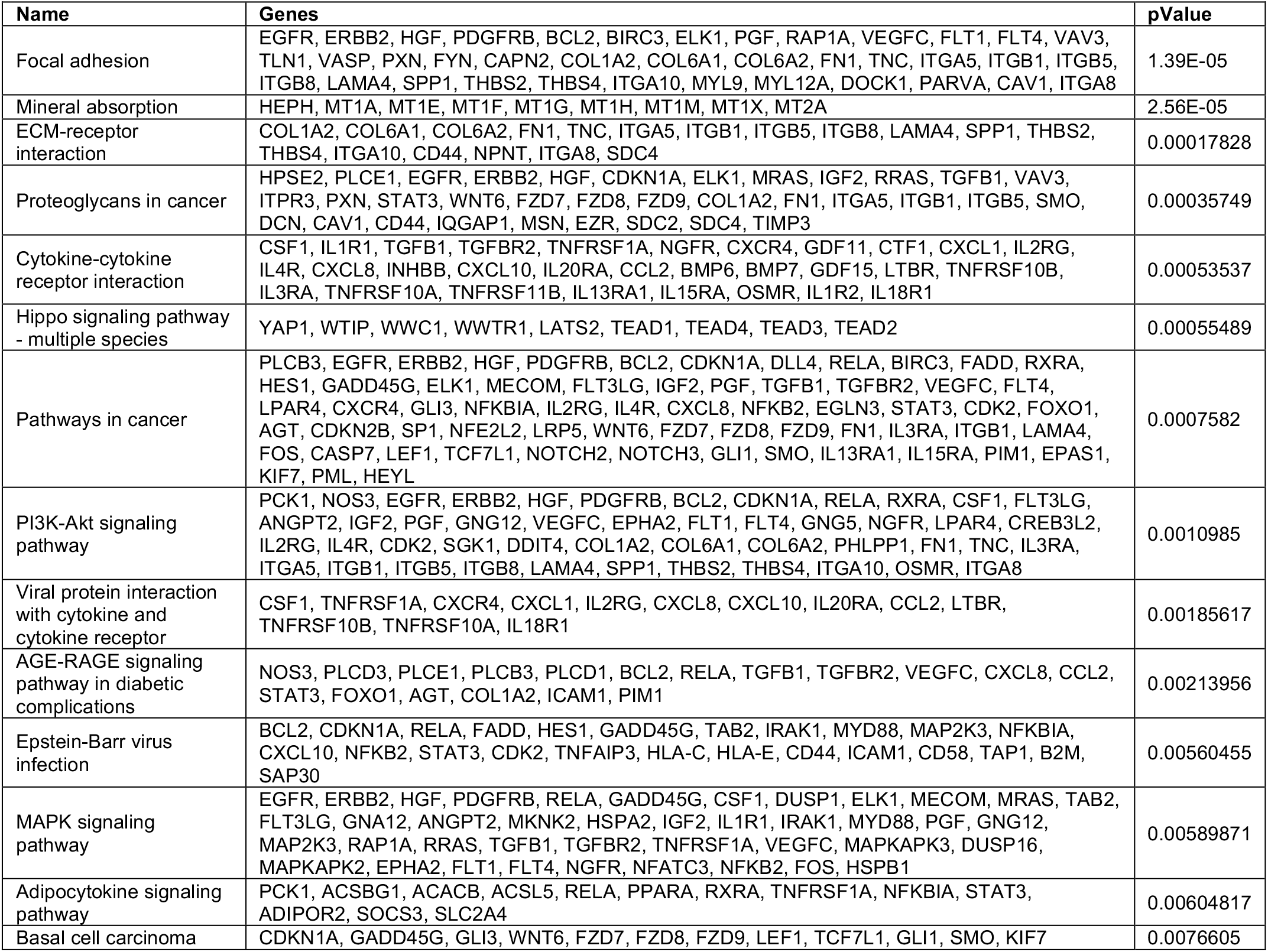
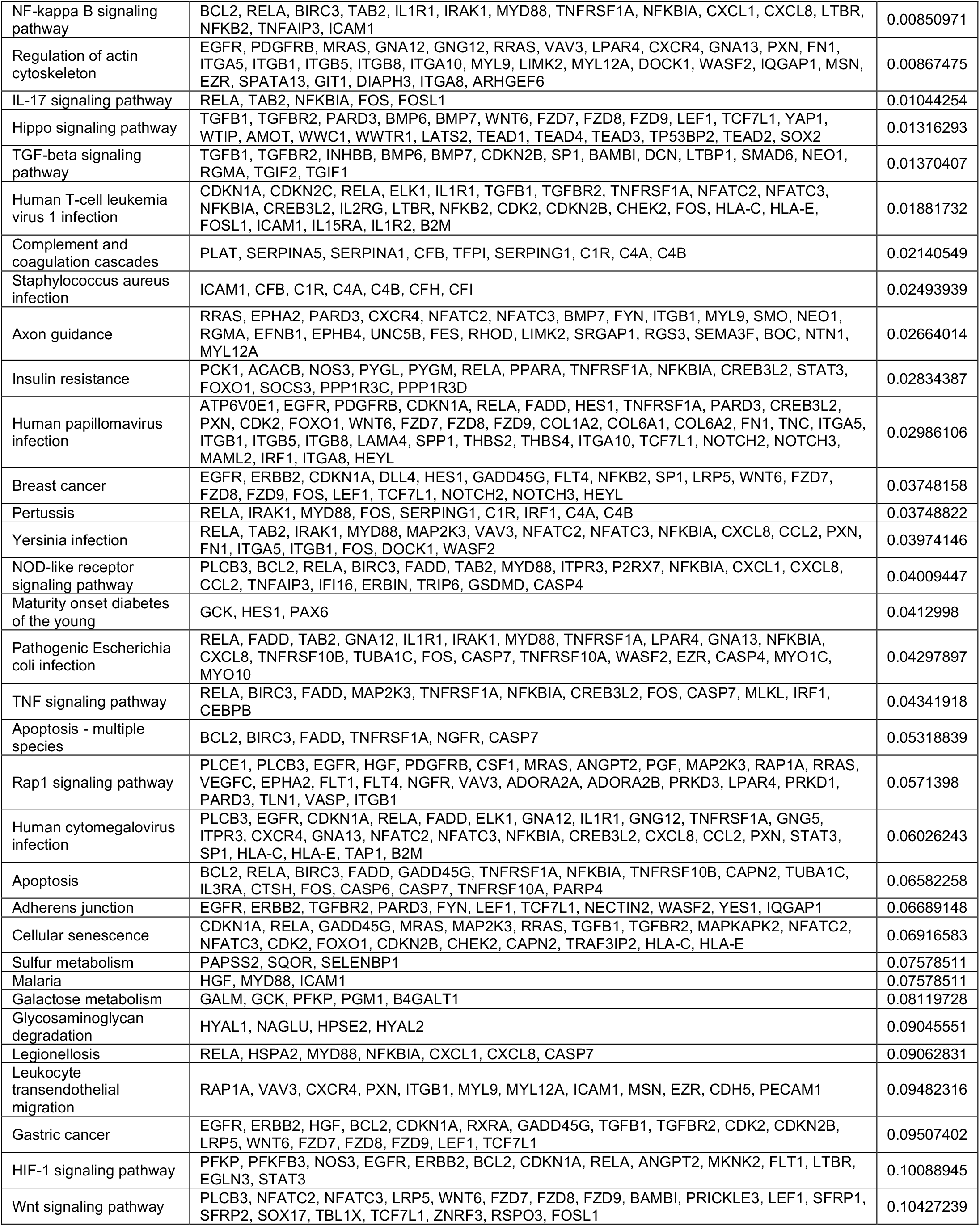
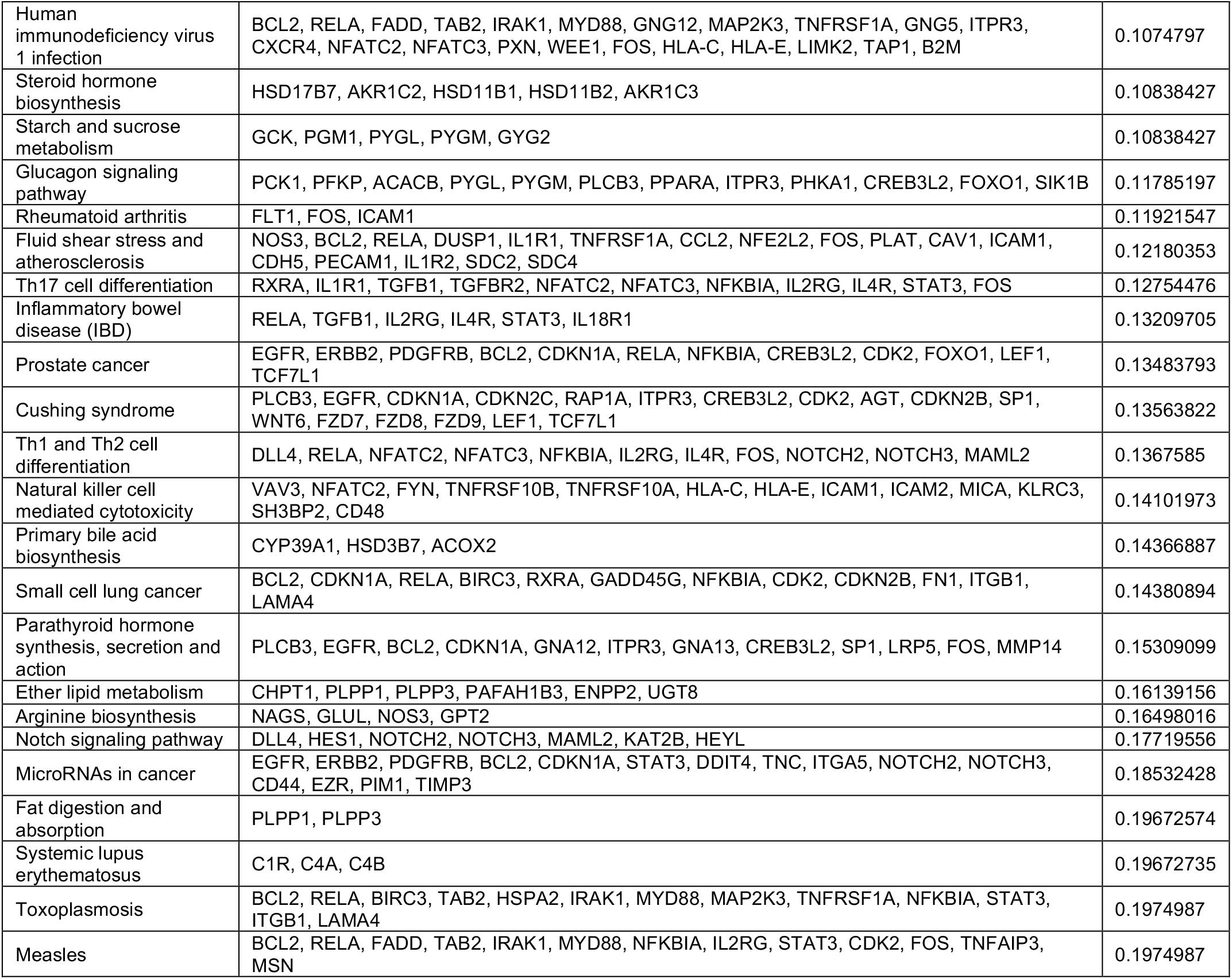
Enriched Kegg signaling pathways using up-regulated genes.

**ST2:** The 61 enriched KEGG signaling pathways.

ST2: The 61 enriched KEGG signaling pathways.
| Name | p-value | Name | p-value |
| --- | --- | --- | --- |
| Focal adhesion | 1.39E-05 | TNF signaling pathway | 0.0434 |
| Mineral absorption | 2.56E-05 | Apoptosis - multiple species | 0.0532 |
| ECM-receptor interaction | 0.0002 | Rap1 signaling pathway | 0.0571 |
| Proteoglycans in cancer | 0.0004 | Human cytomegalovirus infection | 0.0603 |
| Cytokine-cytokine receptor interaction | 0.0005 | Apoptosis | 0.0658 |
| Hippo signaling pathway - multiple species | 0.0006 | Adherens junction | 0.0669 |
| Pathways in cancer | 0.0008 | Cellular senescence | 0.0692 |
| PI3K-Akt signaling pathway | 0.0011 | Sulfur metabolism | 0.0758 |
| Viral protein interaction with cytokine and cytokine receptor | 0.0019 | Malaria | 0.0758 |
| AGE-RAGE signaling pathway in diabetic complications | 0.0021 | Galactose metabolism | 0.0812 |
| Epstein-Barr virus infection | 0.0056 | Glycosaminoglycan degradation | 0.0905 |
| MAPK signaling pathway | 0.0059 | Legionellosis | 0.0906 |
| Adipocytokine signaling pathway | 0.0060 | Leukocyte transendothelial migration | 0.0948 |
| Basal cell carcinoma | 0.0077 | Gastric cancer | 0.0951 |
| NF-kappa B signaling pathway | 0.0085 | HIF-1 signaling pathway | 0.1009 |
| Regulation of actin cytoskeleton | 0.0087 | Wnt signaling pathway | 0.1043 |
| IL-17 signaling pathway | 0.0104 | Human immunodeficiency virus 1 infection | 0.1075 |
| Hippo signaling pathway | 0.0132 | Steroid hormone biosynthesis | 0.1084 |
| TGF-beta signaling pathway | 0.0137 | Starch and sucrose metabolism | 0.1084 |
| Human T-cell leukemia virus 1 infection | 0.0188 | Glucagon signaling pathway | 0.1179 |
| Complement and coagulation cascades | 0.0214 | Rheumatoid arthritis | 0.1192 |
| Staphylococcus aureus infection | 0.0249 | Fluid shear stress and atherosclerosis | 0.1218 |
| Axon guidance | 0.0266 | Th17 cell differentiation | 0.1275 |
| Insulin resistance | 0.0283 | Inflammatory bowel disease (IBD) | 0.1321 |
| Human papillomavirus infection | 0.0299 | Prostate cancer | 0.1348 |
| Breast cancer | 0.0375 | Cushing syndrome | 0.1356 |
| Pertussis | 0.0375 | Th1 and Th2 cell differentiation | 0.1368 |
| Yersinia infection | 0.0397 | Natural killer cell mediated cytotoxicity | 0.1410 |
| NOD-like receptor signaling pathway | 0.0401 | Primary bile acid biosynthesis | 0.1437 |
| Maturity onset diabetes of the young | 0.0413 | Small cell lung cancer | 0.1438 |
| Pathogenic Escherichia coli infection | 0.0430 |  |  |

## Notes

### Competing Interest Statement

The authors have declared no competing interest.

